# Empty drops in scRNA-seq uncover the surprising prevalence of sequestered neuropeptide mRNA and pervasive sequencing artifacts

**DOI:** 10.64898/2026.02.13.705850

**Authors:** Gennady Gorin, Linda Goodman

**Affiliations:** Fauna Bio, Emeryville, CA

## Abstract

The empty drops in single-cell sequencing experiments are an underexplored resource. As such, they present a substrate to ask questions orthogonal to standard single-cell sequencing workflows, calibrate statistical models using simple internal controls, and detect technical outliers which would be otherwise challenging to distinguish from real biology. In this case study, we report a relatively simple procedure to detect sequencing artifacts and make recommendations to reduce the risk of erroneous quantifications. In addition, we report the surprising abundance and co-expression of mRNA coding for neuropeptide-related genes in the empty drops, possibly reflecting underlying physiology.

## 1 Introduction

The empty droplets in a single-cell RNA sequencing experiment have relatively simple statistics, and their gene expression approximately follows the Poisson distribution.

This observation accords with the intuition that the molecules observed in empty droplets are independently randomly incorporated from a mixed cell lysate. However, somewhat surprisingly, the Poisson model is insufficient for a group of genes in a recently published dataset surveying the single-cell landscape of the thirteen-lined ground squirrel hypothalamus [1]. These genes are outliers: they have variances, Fano factors (ratios of variance to mean), and maxima far in excess of those predicted by the Poisson model or its extensions. Some are mitochondrial, pointing to organelle contamination from the lysate: their incorporation is nonrandom because the molecules arrive co-packaged, rather than independently. However, the others — 120 non-mitochondrial genes with maxima in excess of 30 over six samples — are more challenging to account for (Figure 1).

**Fig 1.**
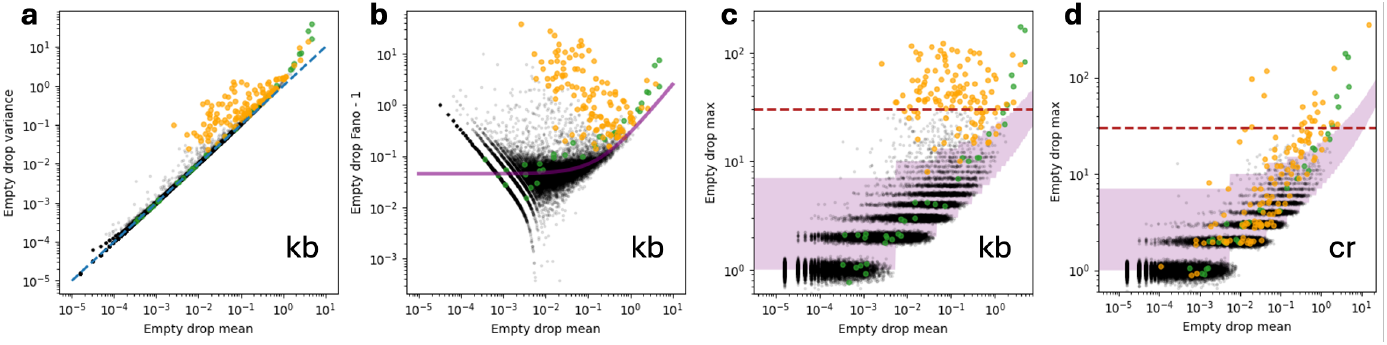
The statistics of empty drops are usually, but not universally, compatible with simple Poisson-like models. **a**. The mean–variance relationship of gene count distributions is near-Poisson, with some exceptions (yellow points: genes of interest; green points: mitochondrial genes; gray points: all other genes; blue line: identity or Poisson). **b**. The mean–Fano factor relationship of gene count distributions can be summarized by a variant of the Poisson model. However, the genes of interest have considerably higher Fano factors than expected (magenta line: theoretical curve given in Equation 3, fit to log-transformed data with mean *>* 10^*−*2^; conventions for points as in **a**). **c**. The mean–maximum relationship of gene count distributions can be approximated by a variant of the Poisson model. However, the genes of interest have considerably higher sample maxima than expected (magenta region: plausible region for the maximum, located between the Poisson lower limit and the Neyman type A upper limit given in Equation 17; dashed line: maximum = 30; conventions for points as in **a**). **d**. The statistical behaviors of the genes of interest are sensitive to the quantification procedure, and Cell Ranger alignment does not produce the pervasive outliers that characterize the *kallisto* | *bustools* results shown in **c** (conventions for points as in **c**).

These outliers may be explained by a staggering variety of possible model violations. For instance, some very small cells may be erroneously assigned as empty droplets; non-mitochondrial RNA may be co-packaged by an analogous mechanism; transcripts with numerous poly(A) stretches may be overrepresented due to multiple priming [2]. However, the majority of these observations appear to be alignment artifacts: many, but not all, outliers that appear in *kallisto* | *bustools* (`kb`) pseudoalignments disappear upon realignment with Cell Ranger. In this manuscript, we attempt to explain these outliers.

## 2 Results

Upon closer inspection, many of the gene quantifications are dominated by artifactual reads. In Figure 2a, we summarize all reads which are 1. assigned to one of the 120 genes of interest by `kb`, 2. *not* assigned to any gene by Cell Ranger, and 3. associated with an empty droplet that has more than two UMIs of that gene. This approach allows us to investigate differences between the two quantification procedures, focusing on the most drastic outliers. Some two thirds of these genes have read pools that closely resemble standard 10X Genomics primers, predominantly concatenations of the template switch oligo (TSO), the feature barcoding capture sequence, the feature barcoding cell barcode, and miscellaneous poly(A) sequences.

**Fig 2.**
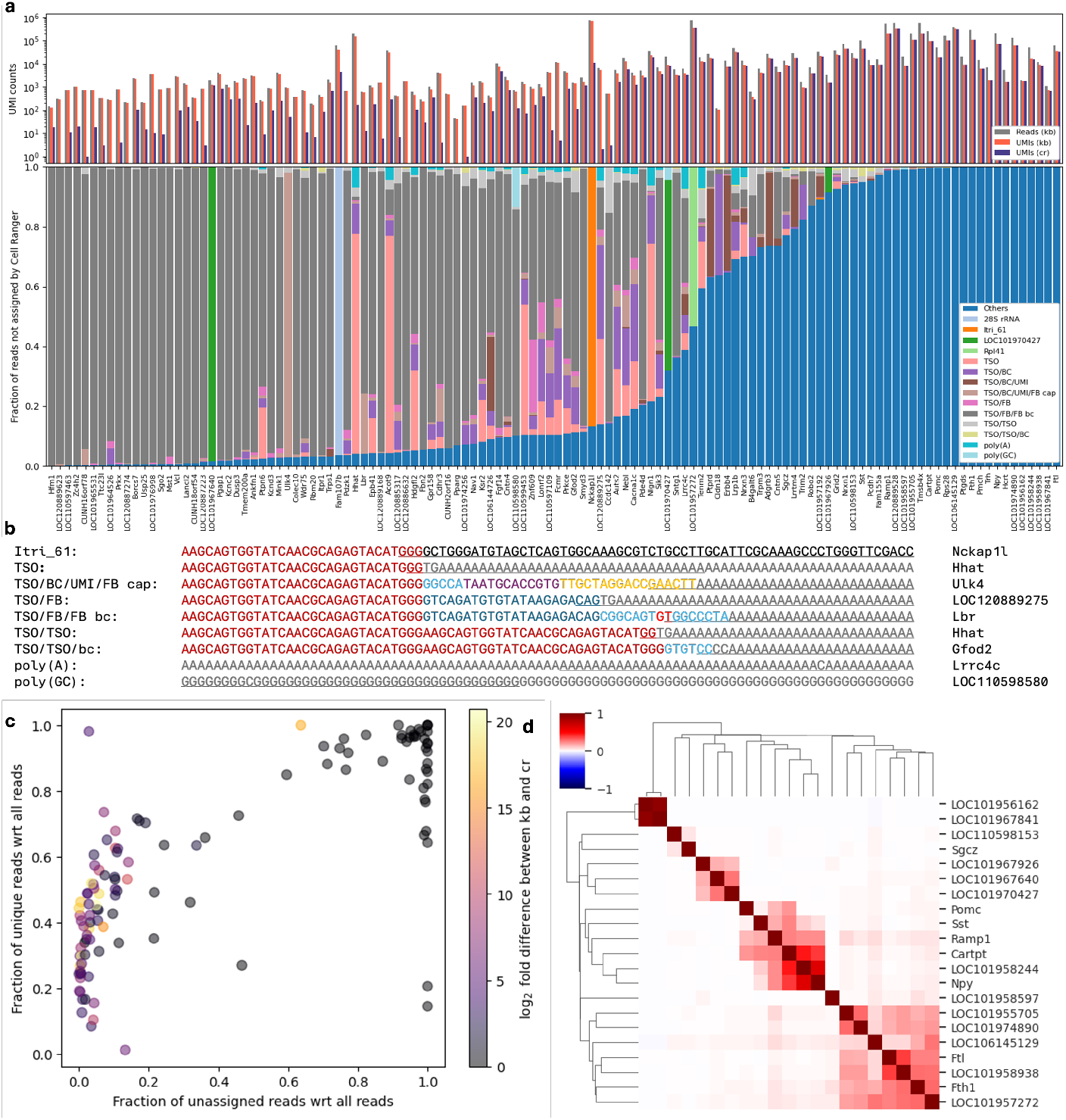
The characterization of outliers. **a**. Top: counts of `kb` reads, `kb` UMIs, and Cell Ranger UMIs across the pool of analyzed reads. Bottom: distribution of read sequence classes. **b**. Example read artifact types, artifacts, and genes (red: TSO; gray: poly(A/G); light blue: cell barcode; bright red: feature barcoding substitution in the barcode; dark blue: feature barcoding 5’ primer; yellow: feature barcoding 3’ capture sequence; purple: UMI, underlining: artifact region that uniquely maps to gene). **c**. Relationship between the diversity of reads and the abundance of reads not assigned to one of the artifact classes (color: quantification difference between `kb` and Cell Ranger). **d**. Correlation-based clustering of the genes most robustly co-vary across samples, relative to theoretical model prediction.

Feature barcoding (FB) is the commercialized version of cellular indexing of transcriptomes and epitopes by sequencing (CITE-seq) [3]. The 10X 3’ v3 single-cell gene expression beads deliver the standard gene expression oligonucleotides, with a sequencing adapter, a cell barcode, a UMI, and a poly(dT) sequence intended to capture poly(A) tails of processed molecules [4, 5]. The same beads bear feature barcoding oligonucleotides, with a different sequencing adapter, a slightly modified cell barcode (with the middle two bases complemented), a UMI, and a capture sequence intended to target complementary sequences conjugated to antibodies [5, 6].

In the absence of these antibodies, the feature barcoding oligos should be inert, as they do not have priming regions. However, we hypothesize that some FB oligos have a poly(A) tail in lieu of the UMI and capture sequence. As a consequence, these oligos are captured by the poly(dT) sequence, undergo TSO capping, fail to fragment due to their short length, are sequenced, and pseudoalign to (mostly intronic) poly(A) stretches in the reference transcriptome. The final several bases of the FB cell barcode provide the non-poly(A) information necessary to disambiguate a read and lead to its erroneous quantification. The pool of such faulty FB oligos is large and nearly identical for a given bead, leading to a large number of reads with similar sequence content but distinct UMIs.

It is possible that the oligo artifacts arise from a different source, but no other chemical mechanism appears to be plausible. For instance, we may hypothesize that the FB cell barcode hybridizes with the gene expression barcode by a strand invasion mechanism [7]. However, it is not clear how such a chimera would produce a cDNA molecule with two barcode sequences.

The artifactual nature of these reads is detectable by other statistics. Reads that originate from endogenous mRNA tend to have high sequence diversity, as the cDNA insert is sequenced from the non-deterministic fragmentation site. Reads that originate from primers have low sequence diversity, as fragmentation does not occur, and the artifactual sequences are overrepresented. The former have similar quantification levels between `kb` and Cell Ranger; the latter are usually discarded by the alignment software but quantified in pseudoalignment.

These trends have some interesting exceptions and caveats.

- *Ulk4* is artifactual, but has high sequence diversity because its reads contain random 12-base UMIs from the FB oligos.
- *Cldn18* is artifactual, but has high sequence diversity because its reads consist of the TSO, a truncated cell barcode, a poly(A) region, and various sequence content of no clear origin. *Lrrc4c* shows a similar structure, but with a highly variable region in lieu of the barcode.
- *LOC106144726* is largely artifactual, but has high sequence diversity because its reads consist of the TSO, cell barcode, and UMI, followed by a poly(T) tail.

The unassigned sequences, which had an edit distance of ≥ 15 to any of the potential artifacts, are a combination of physiological reads which were not aligned by Cell Ranger for one reason or another, sometimes with low sequence diversity and high TSO content; physiological reads which were improperly disambiguated (e.g., using a part of the TSO sequence to assign the gene for a multimapping region); artifacts with no clear source (e.g., a TSO/FB/FB barcode fusion that does not match the cell barcode sequence); and miscellaneous reads that may or may not be artifactual.

These miscellaneous reads urge further consideration: even if their quantifications are artifactual, the reads are real, and their source may reveal interesting and heretofore unrecognized sources of signal or error. In other words: if we observe an unexpected gene hundreds of thousands of times, it is helpful to attempt to understand and account for these data.

*Nckap1l*, which has nearly a million UMIs in the `kb` quantification and two orders of magnitude fewer in Cell Ranger, consists of a pool of nearly identical reads with a TSO followed by a 61-base sequence. This sequence occurs in five other genes in the reference; the disambiguation (and erroneous unique assignment) is based on the last three bases of the TSO. The lack of sequence diversity would suggest the source transcript is short or technical. However, it does not appear to match any known adapter, and BLAST suggests that this sequence is specific to Sciuridae (the squirrels) [8].

We can disambiguate the sequence further by examining the small subset of reads with truncated TSOs. This reveals that the 3’ end of the 61-mer is followed by CCCAGTTCC, then a poly(A)-rich region whose precise sequence is uncertain due to well-known degradation of performance at homopolymers and near sequence ends [9]. This sequence *is* represented in numerous rodent transcripts, sense to *Slc25a30, Riok3*, and *Cdon*, and antisense to *Rtl10, Carnmt1*, and *Pmpcb*, among many others, though no annotated thirteen-lined ground squirrel transcripts are exact matches. Query of raw PacBio sequences used to assemble the 2025 genome reference `mIctTri1` reveals the 61-mer in numerous reads, flanked by `GG` and CCAGTTCC and followed by a poly(A)-rich sequence. These PacBio reads, in turn, partially BLAST to a variety of annotated Sciuridae genes.

The fact that the head begins with `GGG` is suggestive of the antisense TSO priming mechanism [7]: the TSO primes to an internal `GTACCC`; after reverse transcription, the second strand primes by capture of the complementary poly(A) stretch to the synthetic poly(dT). *Carnmt1* meets these criteria, but its sequence has some deviations from the observed reads, and other plausible candidates exist. Although it appears that the reads are consistent with a TSO antisense mechanism, the source remains obscure.

An intron of *Fam107b* has a *>* 1 kb region of almost perfect homology to the thirteen-lined ground squirrel 28S ribosomal RNA (*LOC144374300* in `mIctTri1`, but not annotated in the previous version of the reference). Interestingly, in the human genome, this region contains the 40S ribosomal protein SA pseudogene *RPSAP7*. Therefore, *Fam107b* reads may originate from various copies of ribosomal proteins.

*LOC101957272*, one of the highest-expressed genes with very low sequence diversity, is annotated as a miscellaneous long-noncoding RNA (lncRNA). The reads appear to originate from *Rpl41*, the gene coding for 60S ribosomal protein L41. In the `mIctTri1` reference, it is merged with the neighboring gene *Pa2g4*.

Numerous reads assigned to *LOC101967640* (oxytocin-neurophysin 1) by `kb` more closely match *LOC101970427* (vasopressin-neurophysin 2-copeptin-like), as reported in `mIctTri1`. This is explained by reference limitations. This pair of neighboring antisense genes is homologous to the *OXT* /*AVP* pair in other mammalian genomes. The `HiC_Itri_2` and `mIctTri1` versions of *LOC101967640* are nearly identical. However, the `HiC_Itri_2` version of *LOC101970427* truncates exon 2 and predicts a distant exon 3 (with an intron covering *LOC101967640*). Therefore, numerous reads that unambiguously span the exon 2–3 junction of *LOC101970427* are discarded by Cell Ranger and mapped to a homologous 200-base region of *LOC101967640* by `kb`.

It is somewhat curious to note that some of the overdispersed genes are *co*-expressed, suggesting that their incorporation mechanisms are coupled. The determination of statistical models for covariances is outside the scope of the current investigation, but proceeding heuristically, we find that high deviations between observed and predicted covariances are consistently observed for a subset of genes. These genes fall into several modules, with those of most interest listed below.

- The neurophysins *LOC101967640, LOC101970427*, and *LOC101967926* (oxytocin-neurophysin 1, albeit with a different 3’ UTR).
- The appetite-controlling hormones *Npy, LOC101958244* (*Agrp*), *Cartpt*, and *Pomc*, where *Pomc* is co-expressed with *Cartpt* but no others (reflecting their co-expression in neurons). These are often observed with *Sst*, which codes for a peptide hormone, and *Ramp1*, which modifies neuropeptide activity.
- The hemoglobins, *LOC101956162* (hemoglobin subunit alpha) and *LOC101967841* (hemoglobin subunit beta-S/F-like).
- The ferritins, *Fth1, Ftl*, and *LOC101958938* (ferritin light chain-like). These are often observed with *LOC101957272* (*Rpl41*).
- More weakly, *Sgcz* and *LOC110598153* (apparently a fragment of *Tenm3*).

The co-expression of hemoglobin subunits is somewhat self-explanatory and likely reflects residual RNA in erythrocytes (as in [10]). The ferritin module appears to contain fairly generic neuronal or glial debris.

However, surprisingly, the hypothalamic hormones are strongly overrepresented and co-observed in a way that reflects their cell type distribution (as in Figure 5). It is possible that these observations are derived from miscellaneous debris or dying neurons. However, the limited correlation with standard markers of cell function is evocative of sequestration into vesicles or intracellular granules. In other words, the data may partially reflect the physiological packaging of mRNA coding for hormones and hormone-related genes.

## 3 Discussion

Once they are characterized, the artifacts are trivial to address. For instance, it is straightforward to exclude all reads that contain problematic k-mers from quantification [11]. If we elide reads that contain long poly(A) stretches and the TSO/FB fusion, we retain the advantages of pseudoalignment while reducing the risk of false positives. Repeating the analysis in this way reduces the number of outliers from 120 to 53, predominantly eliminating the FB-related artifacts (Figure 4).

Adapter trimming would not be helpful here, as the artifactual reads originate from the variable barcode region. Poly(A) trimming is more appropriate. Read filtering, e.g., discarding all reads with TSO sequences, creates the risk of false negatives: for instance, in a somewhat extreme case, *LOC106145129* — a fragment of *Lto1* — has numerous reads with *two* TSO sequences followed by a uniquely mapping poly(G)-heavy region, and excluding them may not be appropriate.

The stringency of the proposed filtering procedure is relatively severe. For instance, it eliminates the *Nckap1l* reads, which incidentally contain the excluded 31-mer consisting of the TSO followed by G. The determination of appropriate criteria is a challenging problem and the solution is not self-evident, as these hyperparameters can be tuned in a variety of ways with competing trade-offs.

These artifacts are interesting because they can produce a large number of reads with distinct UMIs even in empty drops, which have few tissue-derived mRNA molecules. Although the artifacts are most straightforward to detect against the background of Poissonian “soup,” the same mechanism may occur in cell-containing droplets, distorting the observed distributions and affecting statistical inference. This risk may be exacerbated by the aggregation of counts across cells (pseudobulking), or mitigated by judiciously setting cutoffs for minimum expressing cell fraction. Relatively simple statistics — such as the overdispersion and maxima in empty drops — can help diagnose these issues as an internal control.

This case study concerns gene quantification in a non-model organism, the thirteen-lined ground squirrel. However, the mechanisms are general; for instance, the TSO/FB/FB barcode artifacts will map whenever an annotated region has a poly(A) stretch adjacent to a short sequence that happens to be partially represented in the barcode pool. As poly(A) stretches are common, this is likely to happen. For instance, the *Hhat* `GGTG`/poly(A) 37-mer occurs in human *IL2RA*; the *Ulk4* 33-mer occurs in *HAGLR*; a 31-base version of the *LOC120889275* `CAGTG`/poly(A) region occurs in *SMOC2*. Therefore, TSO and FB primers can generate unique artifactual alignments even in the absence of barcode sequences. Accounting for barcodes is somewhat more challenging; however, the same risks seem to remain. For instance, a transcript of human *WT1-AS* has a `CACCTGT`/poly(A) 40-mer, and 1,920 10X 3’ v3 barcodes terminate in this heptamer, suggesting these barcodes can give rise to artifactual reads of this gene.

Finally, even after accounting for quantification artifacts, variability between empty drops, and stochasticity in library construction, certain genes *still* show overdispersion, and some are consistently co-encapsulated. The significance of this finding remains obscure. The co-expression patterns may reflect co-condensed cellular debris. Some may be explained by the confinement of RNA pools by residual membranes. Some, such as the capture of 28S ribosomal RNA, may point to the capture of molecular complexes. Others may reflect stabilization by proteins or other RNA species. In addition, reference limitations, such as a single gene being split into two genome entries, can produce inflated sample covariances (Section 4.7.4). Although it is challenging to distinguish between these mechanisms, the enrichment of mRNA coding for common neuropeptides is provocative, and may suggest a physiological mechanism that enriches, sequesters, and releases their transcripts in granules or vesicles — e.g., due to the formation of exosomes for secretion or granules for dendritic transport [12–19] — which proceed to contaminate other droplets in a sequencing experiment.

If this interpretation is correct, the co-expression of neuropeptide RNA is not specific to hibernators and may be detectable in other publicly available single-cell datasets. Single-nucleus data, such as the recently published human HYPOMAP atlas [20], may be less suitable, as the process of isolating nuclei is disruptive. Future studies may fruitfully examine suitably matched mouse data [21, 22]. Ultimately, sequencing generates vast volumes of data and offers numerous avenues of analysis, only a small fraction of which are germane to the topic that motivated its collection. As we demonstrate in this case study, the free release of raw sequences enables reanalyses to ask drastically different questions by leveraging other facets of the data — empty drops, unspliced reads [23], viral sequences [24], ubiquitous long non-coding RNA [25, 26], and many others — and draw attention to pernicious artifacts or reveal otherwise hidden biology.

## 4 Methods

### 4.1 Quantification

We quantified reads using the 2021 `GCF_016881025.1_HiC_Itri_2` (RefSeq `HiC_Itri_2`) genome for the thirteen-lined ground squirrel.

#### 4.1.1 Pseudoalignment

Data were pseudoaligned with *kallisto* | *bustools* (`kb`, implemented in the package `kb_python` 0.29.5). To make the reference compatible with *kallisto* | *bustools*, `gene` entries, which had empty `transcript_id` fields, were removed. The reference was built using `kb ref` in the `nac` configuration, using the `--make-unique` parameter. This reference has the identifier `kallisto_nac_refseq_qc`.

The removal of gene entries is non-destructive because these entries are not required by the `gtf` specification.

The `nac` configuration generates a reference that combines *n*ascent, *a*mbiguous, and *c*DNA quantification. Broadly speaking, this approach enables the quantification of spliced and unspliced mRNA.

Additionally, we generated a second reference, which excludes frequent artifactual sequences. The reference was built using the same configuration, but including the argument `--d-list` with the genomic FASTA and a list of sequences to be masked from quantification. All reads that contain any of the k-mers in these sequences are excluded. These sequences include the poly(A) 32-mer, the 57-base TSO/FB primer fusion, and the 50 unique sequences derived from single-base deletions in the TSO/FB fusion. This reference has the identifier `kallisto_nac_refseq_qc_dlist`.

The gene counts were quantified using `kb count` with the parameters `-x 10XV3` to impose the 10X 3’ v3 chemistry, `--workflow nac` to quantify spliced and unspliced reads, `--filter bustools` to identify putative cell-associated barcodes, `--sum=total` to quantify the sum of all (nascent, mature, and ambiguous) UMIs, and `--num` to output the read numbers in the BUS file.

The pseudoalignments produced the following outputs relevant to the analysis:

- UMI count matrices, including
  – Gene names and cell barcodes used to populate the row and column names,
  – Matrices reporting total, nascent, etc. UMI counts for each gene/cell pair in the `mtx` format,
  – All of the above for the filtered dataset, restricted to the putative cell-associated barcodes,

- Intermediate outputs, including
  – The list of quantified transcripts, including processed and unprocessed RNA species,
  – The list of transcript equivalence classes observed during the pseudoalignment process. In other words, each read is compatible with a set of possible transcripts; the list of those sets is defined in the EC list,
  – The unabridged, unsorted output BUS file, containing the UMI, barcode, equivalence class, and read number for each pseudoaligned read.

#### 4.1.2 Alignment

Data were aligned with Cell Ranger 7.2.0. As Cell Ranger has internal quality control procedures, the genome was not modified. The reference was built with `cellranger mkgtf` and `mkref`. The alignments were performed using `cellranger count` with the parameters `--chemistry SC3Pv3` to impose the 10X 3’ v3 chemistry and `--include-introns true` to quantify unspliced reads (the default setting since Cell Ranger 7.0).

The alignments produced the following outputs relevant to the analysis:

- UMI count matrices, including
  – Gene names and cell barcodes used to populate the row and column names,
  – Matrices reporting total UMI counts for each gene/cell pair in the `mtx` format,
  – All of the above for the filtered dataset, restricted to the putative cell-associated barcodes,
- Intermediate outputs, including
  – The alignment BAM file, containing each read from the original dataset, its cell and molecule barcodes, corrected versions thereof, and (if successfully aligned) the gene name.

### 4.2 Identification of empty drops

We loaded the *kallisto* | *bustools* (`kallisto_nac_refseq_qc`) and Cell Ranger datasets and subset them to the set of genes and cell barcodes that appeared in both datasets.

Although the two quantification procedures use the same whitelist, the barcode correction procedures are slightly different. The lists of identified barcodes are modestly larger for the Cell Ranger datasets. However, the differences are typically restricted to barcodes with very low molecule counts, which are excluded in the following steps.

Next, we excluded all barcodes that had *<* 100 UMIs according to the `kb` total quantification.

The total UMI counts in the datasets are distinctively multimodal, with the peaks coarsely corresponding to neurons (*>* 10^4^ UMIs), non-neurons (10^3^–10^4^ UMIs), empty drops (10^2^–10^3^ UMIs), and potential barcode correction errors (*<* 10^2^ UMIs). These categories should be understood as rough order-of-magnitude estimates, and the ranges overlap.

The problem of describing the distribution of gene counts in empty drops is intimately connected to the problem of modeling contamination, based on the intuition that the contamination should affect empty and cell-containing droplets similarly. Different studies have taken different approaches. For instance, *SoupX* [27] uses the lowest-expression barcodes (here, *<* 10^2^ UMIs) to build an estimator for the contamination. However, we use the terminology established by the authors of the *CellBender* package [28]. Based on Section S.2.1 and Exhibit 3 in S.1.1 of the article [28], we hypothesize that reads associated with lowest-expression barcodes — which are observed in tissue-derived datasets as well as mRNA suspensions — are best explained by barcode misassignment.

Although these reads may be representative of the overall landscape of the ambient transcriptome, they would be discarded in any typical quality control procedure. In addition, they may not accurately represent the barcode/UMI relationship, as they are generated by an atypical mechanism. For these reasons, alongside computational facility (the typical number of empty drops is 10^5^, whereas the typical number of potential barcode errors is 10^6^), we exclude them from investigation.

We narrowed down the putative cell-associated barcodes by taking the union of barcodes in the filtered `kb` and Cell Ranger datasets. We defined the set of empty drop barcodes as the complement of this union.

The goal of this procedure is to establish a consensus set of “typical” empty drops. This filter is relatively conservative to avoid the inclusion of, e.g., unusually small cells.

We subset the BUS outputs to the empty drops using the `bustools` command `correct`, using the list of putative empty drop barcodes as the whitelist (`-w`), and imposing the parameter `--nocorrect` to enforce exact matches to the `kb`-corrected barcodes.

Exact matches are enforced because the output BUS file is corrected with respect to the technology whitelist.

We subset the BAM outputs to the empty drops using the 10X Genomics tool `subset-bam` 1.1.0, using the list of putative empty drop barcodes as the whitelist (`-c`). This procedure uses the corrected barcode tag (`CB`) for filtering.

### 4.3 Identification of outlier genes

Although total UMI counts in the empty drops typically have approximately Poissonian statistics, with variance ≈ mean, this model is violated for a small subset of genes.

These genes are *overdispersed* : their sample variance *S*^2^ is substantially higher than the sample mean 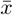; equivalently, the sample Fano factor ^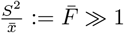^. Furthermore, the observed UMI count maxima are surprisingly high considering the light tails of the Poisson distribution, suggesting that the mechanism generating these data differs from simple contamination by the “soup” of ambient RNA.

Some genes’ unexpected statistics can be explained by known physical mechanisms. For instance, mitochondrial genes (prefixed with `ACI67`) tend to be highly-expressed and overdispersed. As discussed elsewhere [10], this expression pattern is consistent with the encapsulation of entire mitochondria (presumably originating from cellular debris) into empty drops. Once captured, the mitochondria lyse, release their contents, and contribute reads which are ostensibly physiological, but cannot be associated with a specific source cell. Reads from endogenous mitochondria, exogenous encapsulated mitochondria, and exogenous mitochondrial RNA cannot be distinguished, making it challenging to investigate mitochondrial transcriptomics on a single-cell level, but this source of contamination is relatively straightforward to detect in empty drops.

However, non-mitochondrial genes also show overdispersion and high maxima, and their expression is somewhat more challenging to explain from first principles. In addition, some, but not all, of these “outliers” are aligner-dependent: a subset of genes that are overdispersed under *kallisto* | *bustools* quantification show the expected Poissonian statistics under Cell Ranger quantification. Therefore, the expression patterns may be partly attributed to sequence alignment errors. We identified 120 non-mitochondrial genes which had a maximum per-barcode expression of *>* 30 UMIs in at least one of the six datasets aligned by `kb`.

The threshold of 30 UMIs is arbitrary, and intended to detect the most drastic examples of potential artifacts.

To analyze the effect of filtering the *kallisto* | *bustools* index, we used the same list of putative empty drops to analyze the expression quantified with the `kallisto_nac_refseq_qc_dlist` reference.

### 4.4 Read extraction

We identified the transcripts associated with each of the genes of interest based on the transcript-to-gene file generated by `kb ref`. Next, we used the `bustools` command `capture` in the `-s` configuration (transcripts) to identify the BUS entries corresponding to the target transcripts (`-c`) in the empty drops, passing in the corresponding equivalence class (`-e`) and transcript (`-t`) lists.

`bustools capture -s` outputs reads mapping to equivalence classes that *contain*, but are not *constrained to*, the target transcripts. Therefore, the resulting BUS file contains reads that were not quantified, as they map to multiple genes. The output of this step is a superset of the reads of interest.

We decompressed the gene-specific BUS files using `bustools text` using the parameter `-f` to output the read number (BUS flag entry). To narrow down the list to transcripts that were quantified, we filtered the outputs to entries constrained to the target transcripts.

The `nac` configuration of *kallisto* | *bustools* prioritizes exons to break ties and disambiguate reads. In other words, if a read belongs to an equivalence class that maps to an intron of gene A but an exon of gene B, it will be quantified with gene B. We omit these somewhat rare edge cases, and restrict our analysis to reads belonging to equivalence classes that only contain transcripts from a single gene. Therefore, the output of this step is a subset of the reads of interest.

We used the (zero-indexed) read numbers in the BUS files to extract the corresponding (one-indexed) reads from the BAM files. Finally, we used *pysam* 0.22.0 to iterate over the `kb`-quantified reads. For each read, we extracted the original sequence, the corrected cell barcode (`CB`), the uncorrected UMI (`UR`), and the Cell Ranger gene assignment (`GN`).

As we hypothesize the data-generating process in the barcodes with particularly high expression differs from simple Poissonian incorporation, we restrict our analysis to barcodes with *>* 2 counts of each gene of interest, as reported by `kb`. The resulting data, a set of tables reporting the reads underlying the `kb` results for each gene of interest, comprises the substrate for analysis of outliers.

The number of unique cell barcode/UMI pairs in the tables may be compared to the number of UMIs in the count matrices. For `kb`, these counts are typically identical, with the rare exception underestimates induced by exon/intron multimapping (as above). For Cell Ranger, the counts may be higher or lower based on slightly different handling of gene assignment, barcode error correction, and UMI clustering, but tend to be similar.

### 4.5 Assignment of reads

This problem statement is deliberately narrow. It is likely that there are classes of artifacts which are erroneously quantified by both tools; cases where treatment of multimapping leads to different results; artifacts that appear in alignment but do not appear in pseudoalignment; false negatives in one or the other tool. In lieu of being exhaustive, we restrict ourselves to questions of consistency.

We analyzed the reads that were not assigned to any gene by Cell Ranger in an effort to characterize the source of `kb`-specific quantifications.

Reads were assigned to a set of categories, defined using manual curation. We summarized concordance with the following categories of artifactual reads:

- Poly(GC).
  – All bases either G or C.
- Poly(A).
  – All bases A.
- TSO.
  – TSO–poly(A).
  – Variant of the above with the first five bases of the TSO sequence truncated.
- TSO/BC.
  – TSO–gene expression cell barcode–poly(A).
  – Variants of above with the first five bases of the TSO sequence, the first three bases of the barcode, or both truncated.
- TSO/BC/UMI.
  – TSO–gene expression cell barcode–UMI–poly(A).
- TSO/BC/UMI/FB cap.
  – TSO–last five bases of the gene expression cell barcode–UMI–a variant of the FB 3’ capture sequence–poly(A). This case is somewhat unusual and challenging to generalize; the TSO–barcode fusion often contains a poly(G)-heavy region, suggesting aberrant priming. The source of the sequence content that differs from the FB capture sequence is unclear.
- TSO/FB.
  – TSO–FB 5’ capture sequence–poly(A).
  – A variant of the above with the first five bases of the TSO truncated.
- TSO/FB/FB bc.
  – TSO–FB 5’ capture sequence–feature barcoding cell barcode–poly(A).
  – Variants of the above with the first five bases of the TSO or the first three bases of the barcode truncated.
- TSO/TSO.
  – TSO–TSO–poly(A).
  Variants of above with the first five bases of either or both of the TSO sequences truncated.
- TSO/TSO/BC.
  – TSO–TSO–gene expression cell barcode–poly(A).
  Variant of above with the first three bases of the barcode truncated.
- Itri 61.
  – TSO–the squirrel-specific 61-mer observed for *Nckap1l*.

We used the Levenshtein distance to quantify the difference between observed reads and the artifactual sequences. For the first two entries, this was simplified to counting the number of non-GC or non-A bases in each read. When a UMI was present in the hypothetical structure, the location of the UMI-containing region was discarded from the read sequence. Therefore, the Levenshtein distances are, strictly speaking, computed over 91- and 79-base spaces for UMI-containing and non-UMI-containing artifacts, respectively.

Additionally, we aligned each read to a set of candidate sequences using the Needle aligner [29]:

- Thirteen-lined ground squirrel 28S ribosomal RNA, the single-exon transcript *LOC144374300* in `mIctTri1`.
- Canonical transcripts for mouse *Pa2g4* and *Rpl41*.
- Canonical transcripts for `mIctTri1` *LOC101967640* and *LOC101970427*, oxytocin and vasopressin.

The pairwise aligner was run in `local` mode, with gap score set to − 2.5 to discourage non-contiguous alignments. With a slight conflation of terms, we treated the complement of the alignment score (i.e., 91 less the score) as a “distance” between each read and the candidate sequence.

All sequences with distance *>* 15 to all of the candidate sequences or artifacts were assigned to the “Others” category. Sequences with distance ≤ 15 were assigned to the category with the lowest distance.

This analysis is necessarily cursory. The categories of artifactual sequences were selected based on manual inspection, and more systematic approaches are possible. It is likely the fifteen-base threshold can, on one hand, include reads that severely diverge in structure, and on the other exclude plausible alignments (e.g., TSO/*Rpl*41 reads).

### 4.6 Clustering

To understand the co-expression of genes in the empty drops, it is helpful to characterize the co-expression of genes in the cell-containing droplets. We clustered the *scVI* embedding of the Cell Ranger counts [30], restricting analysis to barcodes which were identified as cells in both of the Cell Ranger and *kallisto* | *bustools* analyses.

This set of barcodes is not the complement of set used in the empty droplet analysis, as it excludes the more ambiguous barcodes which were identified as cells by one pipeline but not another.

Although this procedure was relatively coarse (*scVI* 1.0.4, 500 most variable genes, 50 epochs, Leiden clustering with resolution 1.4), it successfully recapitulated the elementary composition of the hypothalamus: neurons that have glutamatergic, GABAergic, and (rarely) histaminergic molecular profiles, and a diversity of glia, predominantly astrocytes and oligodendrocytes. Within the neurons, we observe three distinct subpopulations that express the neurophysins (glutamatergic); the anorexigenic neuropeptides *Cartpt* and *Pomc* (glutamatergic); and the orexigenic neuropeptides *Npy* and *LOC101958244* (*Agrp*), alongside *Cartpt* (GABAergic). The genes *Ramp1* and *Sst* are expressed with low specificity across the latter two subpopulations.

### 4.7 Theory

In this section, we define a probabilistic model and derive its statistical quantities. Most of these identities are closely related to results reported in [23, 31].

#### 4.7.1 Single-gene foundations

First, consider a random variable Λ that is determined by the concentration of primers and reagents across droplets. Λ has mean *µ*_Λ_ and variance 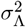.

Next, consider the process of library construction. The distribution of molecule counts of a particular gene g across droplets is given by the random variable *X*_g_, which we assume independent of Λ. The determination of *X*_g_ for all g is the inverse problem of reconstructing gene expression from a noisy single-cell experiment, and is generally complex and underspecified. However, if we restrict analysis to empty drops, we can plausibly hypothesize that 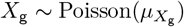. In other words, the “expression” of a given gene reflects simple independent and identically distributed diffusion of molecules into a droplet.

The average 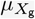 is, in turn, related to the tissue-wide expression of g, although not necessarily the expression of g in the cells actually captured in the experiment. For instance, a single-cell experiment performed on dissociated muscle tissue may fail to capture myofibers, which are too large to encapsulate; however, the dissociation process will nevertheless lyse some myofibers and release their contents, which will to some degree contaminate other cells and empty droplets.

Following [10], we hypothesize that the distribution of (physical) cDNA library molecules across genes g and cells c is drawn from *Y*_gc_ ∼ Poisson(*α*_g_*X*_g_Λ_c_). In other words, the construction of the cDNA library consists of a Poisson birth process that depends on the rate of capture *α*_g_, the number of molecules *X*_g_, and the per-droplet reagent concentration Λ_*c*_. The rate of capture is presumed identical across droplets, encoding the assumption that the chemical environments are similar enough to neglect any differences.

In the *Monod* model [23], Λ is also presumed identical across droplets, producing a Poisson–Poisson, or Neyman type A, distribution of molecules in the cDNA library.

We further hypothesize that the distribution of (bioinformatic) UMIs is described by binomial sampling, such that *Z*_cg_ ∼ Binom(*β*_g_*Y*_gc_). In other words, the number of molecules *actually* observed after amplification, fragmentation, sequencing, and quantification is dependent on the gene identity — depending on the gene, the cDNA molecules may be more or less amenable to amplification, or produce reads that cannot be distinguished due to sequence identity with a different gene. *β*_g_ is identical across droplets as these steps of the sequencing process take place after emulsion breaking.

From Poisson thinning, *Z*_cg_ ∼ Poisson(*α*_g_*β*_g_*X*_g_Λ_c_), allowing us to define the variable *a*_g_ := *α*_g_*β*_g_ that represents the “lumped” and Λ-normalized rate of observing any particular molecule of g in a droplet. In addition, since Λ only appears in the product with *a*_g_, we can set *µ*_Λ_ = 1 (amounting to a unit conversion). In other words, under this model, we may assert *Z*_gc_ ∼ Poisson(*a*_g_*X*_g_Λ_c_).

Consequently, the expectations across droplets 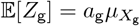 and

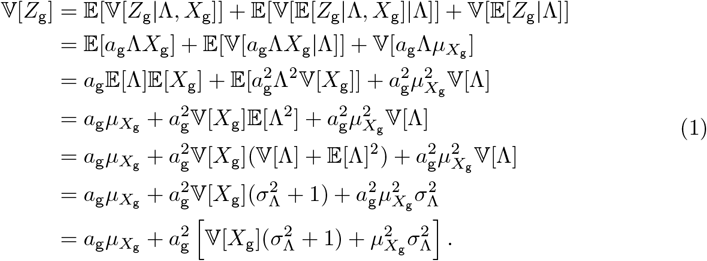

This holds regardless of the distribution of the underlying *X*_g_ variable. Now, assuming the Poisson form 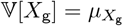, we find

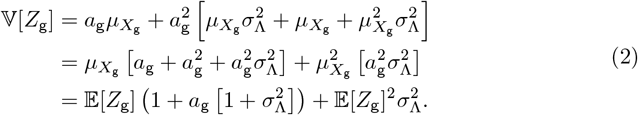

Additionally, the theoretical Fano factor 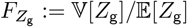 is

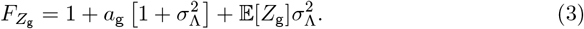

This identity is primarily interesting because it predicts scaling behaviors slightly different from simpler models. Under a zeroth-order approximation, *Z*_g_ is Poisson and 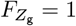. Under a first-order approximation, *Z*_g_ is negative binomial and 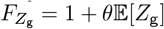 for some dispersion *θ*. In other words, the current model predicts that 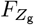 is slightly above unity for low 𝔼 [*Z*_g_], whereas the Poisson and negative binomial model suggest that this limit is unity. If we use the Neyman type A limit 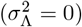, the Fano factor is constant at 1 + *a*_g_.

The functional form of the full distribution is somewhat unwieldy. However, the Neyman type A probability-generating function (PGF) for a single gene, conditional on Λ, is:

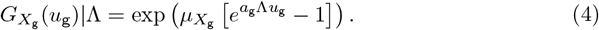

This distribution is infinitely divisible for a constant *a*_g_Λ. In other words, if two molecular species g_1_ and g_2_ have 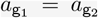, they can be conflated, and their counts can be added, yielding another Neyman type A distribution with parameters. However, this fails to hold if 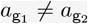.

Even for a single gene, different molecular species —– nascent RNA in the process of transcription and various isoforms — may well have different *a*_g_, e.g., due to varying numbers of poly(A) regions, stability, or bioinformatic identifiability. Nevertheless, this approximation appears to be sufficient to capture multiple interesting features of the data.

The Neyman type A distribution affords useful limiting cases, which can be obtained by Taylor expansion with respect to the parameters. Keeping 𝔼 [*Z*_g_] constant,

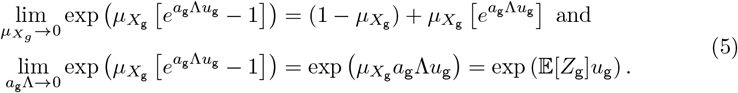

In other words, when the true expression is low, the distribution becomes a Poisson–Bernoulli mixture, i.e., a zero-inflated Poisson distribution: if *X*_g_ = 0, zero UMIs are observed; if *X*_g_ = 1, a Poisson sample of UMIs is generated. On the other hand, when the sampling rate is low, the sequencing becomes Bernoulli, and the overall distribution degenerates to a Poisson with indistinguishable parameters.

#### 4.7.2 Fano factor curve estimation

We assume that *σ*_Λ_ is constant across genes, but *a*_g_ is a gene-specific capture parameter. Nevertheless, we *can* simply fit Equation 3 as though *a*_g_ = *a* across all g, keeping in mind that this assumption is unlikely to hold in practice. The results are illustrated in Figure 1b by a magenta curve: we restrict analysis to non-”interesting,” non-mitochondrial genes with mean *>* 10^*−*2^ in the empty drops and fit the curve 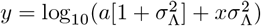, where *y* is the log_10_ of the sample Fano less 1, and *x* is the sample mean. Despite the simplistic approach, the fits appear to recapitulate the shape of the curve, with the exception of the shallowly sequenced dataset SRR25918349.

Intuitively, we may expect the distribution shape obtained by jointly fitting the sample statistics of various genes to correlate with the variability in the empirical “cell sizes,” i.e., the total UMI counts with all gene identity information discarded. As shown in Figure 3a (using Cell Ranger quantifications), this intuition is largely correct: the inferred values of *σ*_Λ_ are generally proportional to the coefficient of variation of the UMI counts, although the inferred value for SRR25918349 are higher than expected. The constant of proportionality is approximately 1.4.

The coefficient of variation (ratio between standard deviation and mean) is chosen because the Λ random variable is defined to have unity mean.

The quantity ^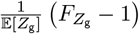^ is named the overdispersion in [31]. Its limit as 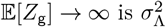. The values reported here (0.15–0.54, with 0.94 for the outlier) are in qualitative, but imperfect agreement with those reported in Figs. 4b and 5e of [31], which may speak to methodological or chemical differences.

As shown in Figure 3b, there is no clear relationship between, e.g., the inferred *a* value and the average of the total UMI count, as the latter is confounded by the overall expression levels. The *a* values range from 0.006 to 0.021, i.e., the ratio of molecules to UMIs generated from them is on the order of 100 : 1. This is surprisingly low, as even the first version of the technology claimed 2–25% conversion [4]. In addition, it is quite a bit lower than the values obtained by jointly fitting nascent and mature count distributions while assuming 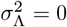, approximately 10% [23]. Reconciling these discrepancies will likely require more sophisticated modeling.

The inference is coarse, and can be improved by appropriately accounting for uncertainty in the Fano and mean estimates (e.g., by bootstrapping). Additionally, the curve can be refined by selecting the most “Poisson-like” genes [31], at the risk of introducing confirmation bias.

#### 4.7.3 Multiple genes

Now, consider the case of two genes g_1_ and g_2_, whose molecule count variables 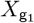 and 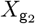 are independent. From the law of total covariance, it follows that

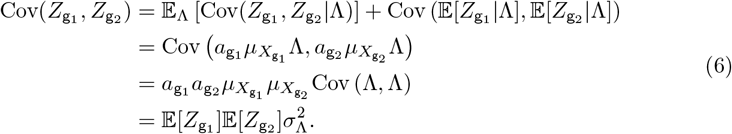

In other words, even in the absence of systematic gene–gene coexpression in the empty drop “soup,” some covariance will nevertheless be observed due to the variability in *Z*. This holds regardless of the distribution of the underlying 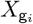 variables, so long as they are independent.

Although this expression looks deceptively similar to Equation 3 — it reports a moment prediction which can be compared to real data based on some estimate of the mean — it is somewhat less useful: the predictions are strictly positive, but the sample covariances are, in a lot of cases, negative (cf. pooled RNA results in [31]). Simple bootstrapping confirms that the estimates are unstable.

Nevertheless, we can use the plug-in estimate of 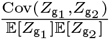 for each pair of genes and identify the pairs of genes that *consistently* display high normalized covariances (in the top quintile). Here, we restrict the analysis to the 120 genes of interest, but use the Cell Ranger data to avoid pseudoalignment artifacts. Twenty-one of these genes have consistently high mutual covariances in all six samples.

A considerable literature exists on the distributions of sample covariances; however, to our knowledge, analytical results are largely restricted to normal distributions [32] or approximate expressions obtained through the central limit theorem. In the domain of low, integer expression, these strategies may not be appropriately calibrated. The statistical significance of high covariance may, in principle, be tested by appropriately bootstrapping; a more statistically rigorous approach may fit multivariate distributions directly. We leave this lacuna to future work.

#### 4.7.4 Single gene, multiple observables

Now, consider the case of a gene subject to the following processes:

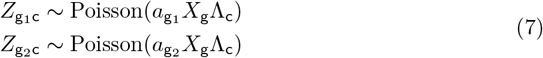

This scenario represents the case where a single gene g can be independently captured at different regions, giving rise to different observed products g_1_ and g_2_.

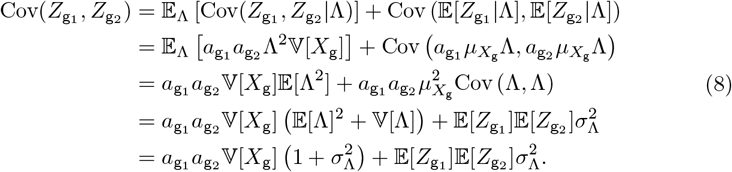

This holds regardless of the distribution of the underlying *X*_g_. If we assume that this random variable has the negative binomial-like variance

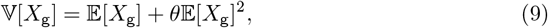

where dispersion *θ* = 0 for the Poisson limiting case, we yield

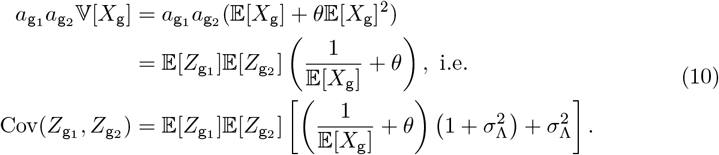

This expression cannot be readily constrained based on data about *Z*: the distribution of underlying molecule counts *X*_g_ is unknown. However, regardless of assumptions regarding its form, the covariance is strictly higher than that given in Equation 6, more so when the average molecule counts are lower or the dispersion is higher.

In the absence of droplet-to-droplet noise in the capture rate (i.e., 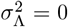), the covariance of g_1_ and g_2_ is simply

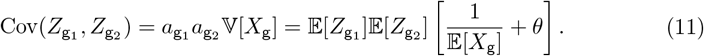

#### 4.7.5 Distribution of order statistics

The distribution of the maximum *M*_*n*_ of *n* observations {*X*_1_, …, *X*_*n*_} drawn from a distribution with cumulative distribution function (CDF) *F*_*X*_ is 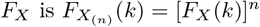.

For a Poisson(*µ*) distribution,

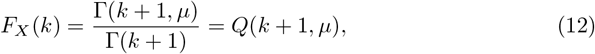

where G(·) is the gamma function, Γ(·, ·) is the (upper) incomplete gamma function, and *Q*(·, ·) is the regularized (upper) incomplete gamma function. Then

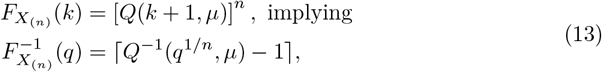

where the inverse is taken with respect to the first argument.

The evaluation of this equation requires somewhat brute-force root finding. However, a result due to Briggs et al. [33] provides an approximate expression for the location of the maximum of *n* realizations of a Poisson(*µ*) random variable:

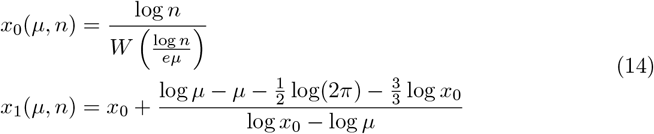

Therefore, *f*_Briggs_(*µ, n*) := ⌈*x*_1_(*µ, n*) ⌉ can be used as a reasonable estimate for the Poisson limit.

There are no comparable results for a Neyman type A distribution. Nevertheless, since this distribution consists of the Poisson being compounded with itself, we can coarsely constrain it:

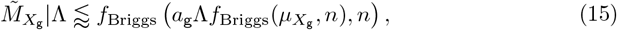

where 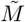 denotes the approximate maximum. In other words: the Neyman type A maximum cannot be computed directly; however, we *can* estimate the maximum of the Poisson molecular distribution, condition on this maximum (scaled appropriately), then obtain the maximum of the UMI molecular distribution. This is an overestimate: we condition, but compute as though the maximum is independently achieved in both realizations of the Poisson draw.

Equation 15 is underconstrained and cannot be compared to data as it stands. However, we can rewrite the expression (for now assuming Λ constant) as

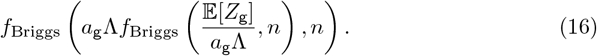

In other words, we note that 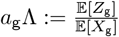, the ratio between average UMIs observed and molecules physically present in the droplet. It appears reasonable to suggest that this fraction should be *<* 1: only a small portion of molecules is captured and sequenced. Therefore, we can tentatively propose that the maximum observed under this model, conditional on Λ constant and the assumption that *a*_g_Λ *<* 1, is

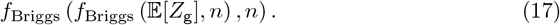

Variability in Λ may increase the maximum. However, we choose to neglect this. Although a procedure similar to the above can be performed using some estimate of Λ(*n*), it is more questionable: assuming that the maxima of the underlying molecule distribution *and* of the Λ distribution *and* of the sequencing distribution are achieved in a single droplet, simultaneously, appears implausible. In addition, the range of per-barcode UMI totals, potentially a proxy for Λ, is relatively narrow. Outliers exist, but the 99th percentile is some 1.5 − 3× higher than the average. As the dependence of *f*_Briggs_ upon the *µ* parameter is very weak, this correction appears unlikely to make a major difference on a per-gene basis.

The assumption that *a*_g_Λ *<* 1 may well be violated: for example, a droplet may have a small number of very long, poly(A)-heavy transcripts which get sequenced numerous times. However, this is a somewhat unusual case, and worth characterizing.

## Data and code availability

The intermediate data, including AnnData objects, per-gene BUS and BAM files, and read annotations, are provided at https://zenodo.org/records/18473667. The scripts and notebooks used to analyze the data are provided at https://github.com/Fauna-Bio/GG_2026.

## Acknowledgments

L.G. is a co-founder and CTO of Fauna Bio. G.G. is an employee of Fauna Bio. The authors would like to acknowledge Delaney Sullivan, Phil McNamara, Meichen Fang, and Tara Chari for their assistance in the preparation of this manuscript.

## Declaration of interests

L.G. is a co-founder and CTO of Fauna Bio. G.G. is an employee of Fauna Bio.

## Supporting figures

**Fig 3.**
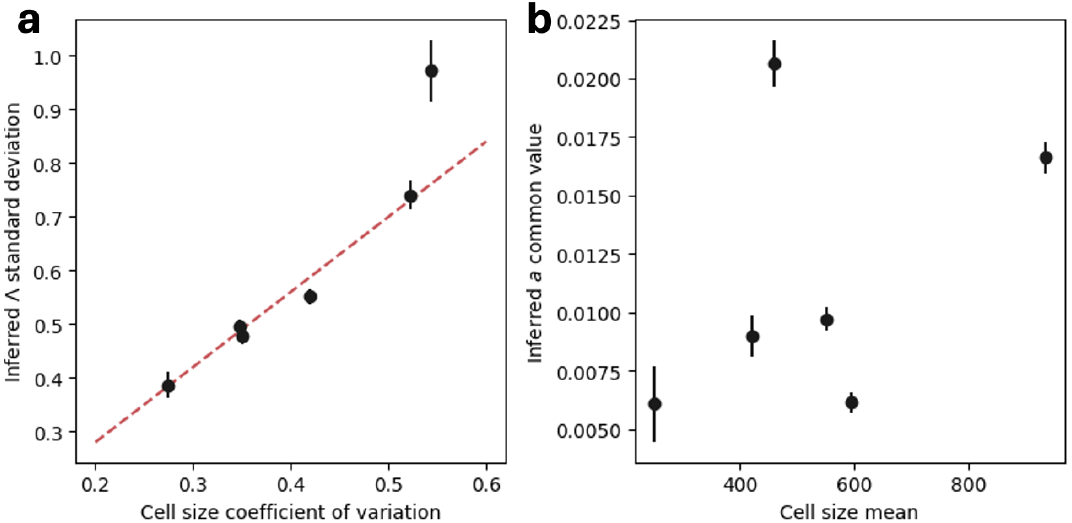
The relationship between the parameters of Fano/mean curve (e.g., magenta line in Figure 1b) and summary statistics of the total UMI count distribution for six datasets (points: datasets; error bars: approximate 95% uncertainty estimates obtained from the parameter covariance matrix). **a**. Relationship between inferred *σ*_Λ_ and total UMI coefficient of variation (dashed line: proportionality at *y* = 1.4*x*, shown to guide the eye). **b**. Relationship between inferred *a* and total UMI average.

**Fig 4.**
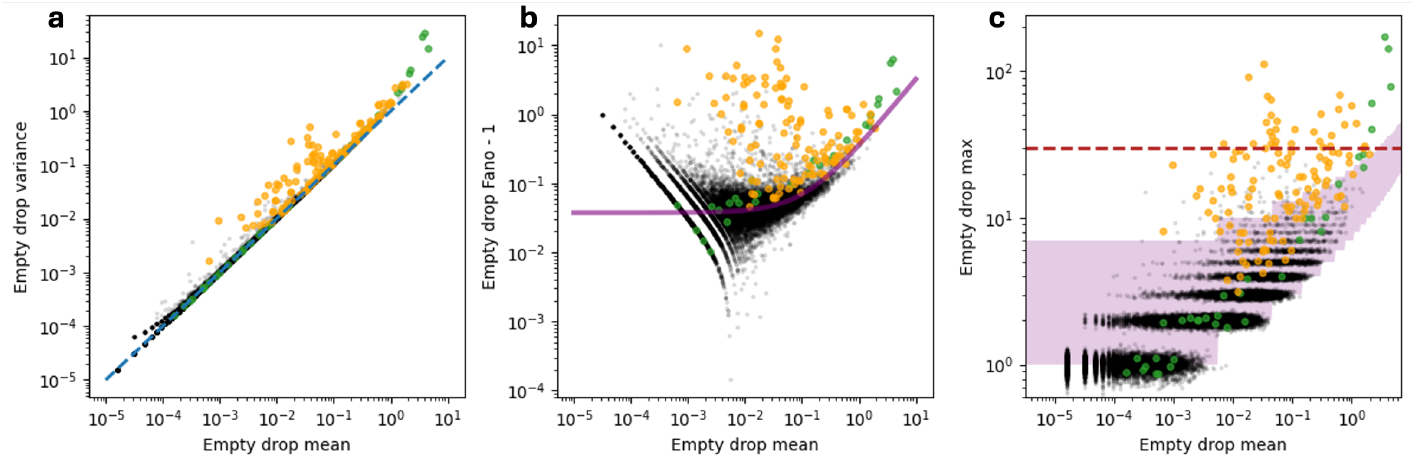
Figure 1a–c reproduced with a filtered `kb` index. The exclusion of known artifactual sequences reduces — but does not eliminate — the presence of outliers.

**Fig 5.**
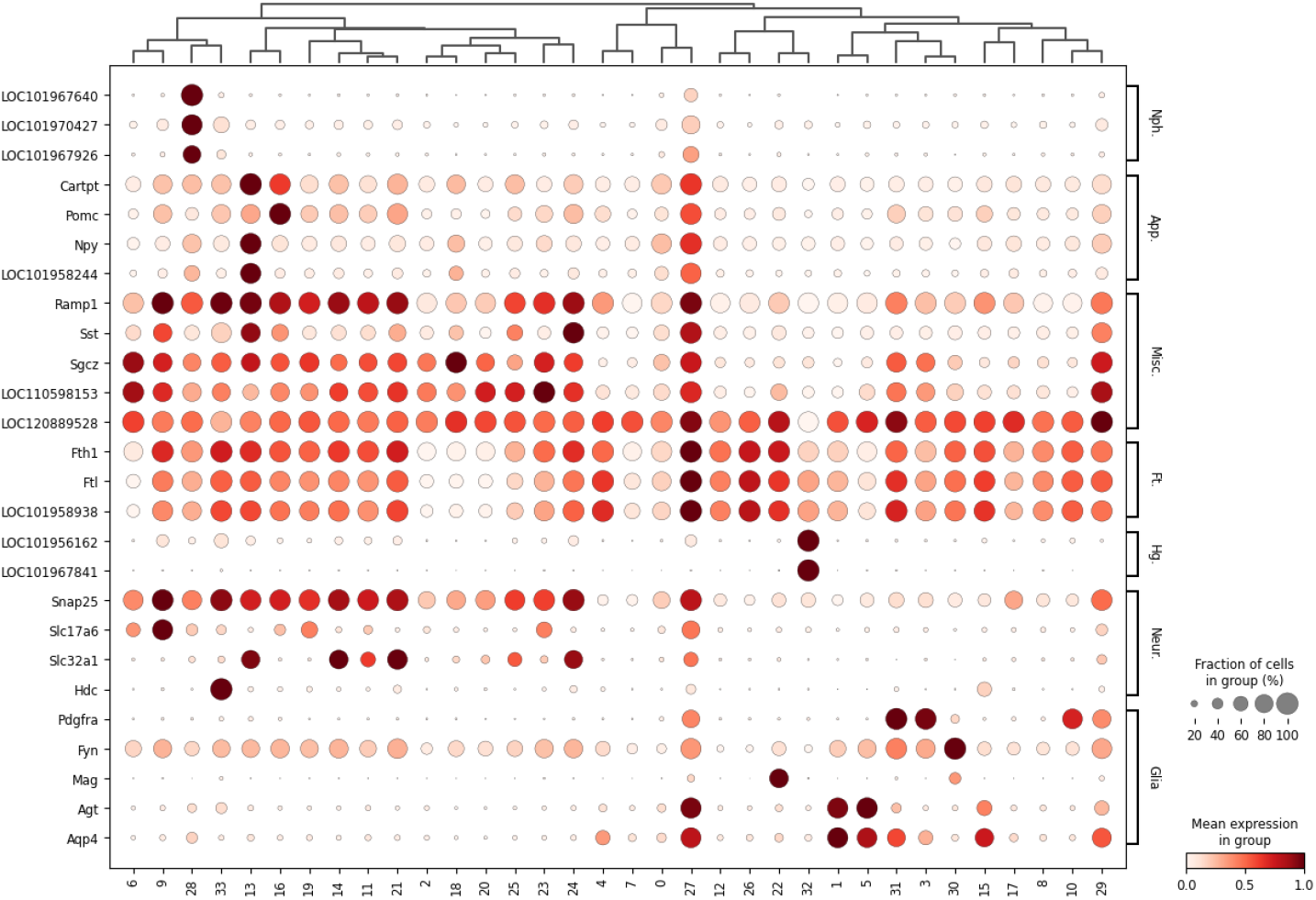
A broad overview of the clusters observed in the Mohr et al. data [1] (Nph.: neurophysins; App.: appetite-controlling hormones; Misc.: miscellaneous genes of interest; Ft.: ferritins; Hg.: hemoglobins; Neur.: neuron markers; Glia: a selection of glial markers.

## Notes

https://github.com/Fauna-Bio/GG_2026

https://zenodo.org/records/18473667

